# Spatio–temporal modelling of *in vitro* influenza A virus infection: the impact of defective interfering particles on type I interferon response

**DOI:** 10.1101/2025.10.09.681519

**Authors:** Yimei Li, Bjarke Frost Nielsen, Simon A. Levin, Aartjan J.W. te Velthuis, Bryan T. Grenfell

**Author notes:** Corresponding authors (AJWTV), (BTG).

## Abstract

Defective interfering particles (DIPs) are incomplete viral genomes that modulate infection by competing with wild–type viruses and activating innate immunity. How DIPs interact with type I interferon (IFN) in spatially structured environments remains unclear. Focusing initially on influenza A, we developed a spatially explicit, stochastic model of *in vitro* viral infection integrating virus and DIP replication, IFN signalling, and alternative dispersal modes. We find that: (1) our model captures the ring–like and patchy plaque morphologies observed experimentally; (2) IFN production peaks at an intermediate DIP ratio, reflecting a trade–off between early immune activation and sufficient co–infection; and (3) even a small fraction of long–range spread of virus and DIPs escape containment despite longer IFN ranges; this causes stronger antiviral responses but earlier peaks in lysis at similar levels of cell loss. The model is available as an interactive platform: https://shiny-spatial-infection-app-production.up.railway.app/.

## 1 Introduction

Populations of RNA viruses, such as influenza A virus (IAV), are generally composed of both standard and nonstandard viral particles. Nonstandard particles contain genomes that functionally differ from the wild–type. A special group of nonstandard particles are the defective interfering particles (DIPs) [1, 2, 3]. DIPs are virus–derived particles with incomplete or defective viral genomes (DVGs) that cannot replicate without co–infection with a standard virus. DVGs with a smaller genome size, also called deletion–containing viral genomes (DelVGs), may replicate more quickly and outcompete standard viruses for host cell resources, while altered gene expression can yield proteins that interfere with standard virus protein function [3, 4, 5, 6]. Despite these differences, DIPs co–exist and co–transmit with standard viruses in tissue culture as well as in humans, influencing infection spread and disease outcome by modulating viral adaptation, virulence, and immune evasion [7, 8, 5, 9, 3, 10, 11].

### DIPs and IFN responses

An important way in which DIPs shape infection dynamics—particularly in IAV infection—is by triggering stronger innate immune responses [12, 13]. Replication and transcription products derived from shortened DIP genomes are more potently detected by pattern recognition receptors (PRRs) such as retinoic acid–inducible gene I (RIG–I) than full–length wild type genomes [14, 3, 15]. Activation of these PRRs initiates signalling pathways that lead to expression of type I and III interferons (IFNs) and various pro–inflammatory cytokines, which are important for dendritic cell activation, fever, and shaping adaptive immunity [16], making DIPs critical actors in virus–host interactions [3]. DIPs have been shown to impact disease progression, act as antiviral and antitumour agents, and interfere with the efficacy of live attenuated vaccines (e.g. IAV [17], measles [18], rubella [19], and others [20]). These features have renewed interest in quantitative understanding of how DIPs alter infection dynamics and innate immunity [21, 9].

### Limitations of existing models

Several computational studies have explored within–host *in vitro* infection dynamics, yet critical gaps remain in integrating spatial resolution, immune responses, and viral diversity. Traditional ordinary differential equation (ODE) models, which represent infection and immune processes as changes in average population densities over time, have assumed that IFN reduces viral production by infected cells, decreases the likelihood of infection, or renders uninfected cells refractory to infection [22, 23, 24]. However, such models lack spatial information and implicitly assume uniform IFN protection across the tissue, even in cases where DIPs are included [4, 25]. A spatial framework for *in vitro* infection dynamics was introduced by Howat et al. [26], who developed a stochastic, spatially explicit computational model of IFN responses. However, their model assumed instantaneous and global IFN diffusion. Once triggered, IFN did not decline, and cells that transitioned to the antiviral state remained permanently antiviral. In addition, the assumed delay before this transition was very long (95 hours), and the model did not incorporate DIPs, thereby potentially overlooking gradients, gradual activation dynamics, and DIP–IFN interactions. Other models have examined DIP–virus interactions at intracellular or population levels but without immune components such as IFN signalling, limiting their applicability to immune–competent systems [27, 28, 29, 30, 31]. Akpinar et al. linked single–cell dynamics to plaque–level outcomes using a spatial cellular automaton, showing how spread depends on initial co–infection with viruses and DIPs [9]; Baltes *et al.* [32] directly visualised DIP–virus co–transmission, distinguishing virus–only, DIP–only, and co–infected cells through different fluorescent markers; and Liang *et al.* [33] analysed such plaque images using binarisation and geometric segmentation. While each of these studies highlighted the spatial structure of viral particles spreading, none incorporated IFN responses, leaving open how innate immunity modulates DIP–virus dynamics.

To bridge these gaps, we developed a stochastic, cell–based computational model of *in vitro* infection dynamics, incorporating DIPs, IFN signalling, and spatial structure (see Figure 1 and Figure S1 for a schematic representation). Our framework is designed to explore how DIP–virus interactions generate heterogeneous plaque structures, how different dispersal routes (cell–to–cell spread, finite–rate diffusion, or rare long–range jumps) influence infection dynamics, and how IFN signalling—whether instantaneous or finite–rate diffusion—modulates immune protection. Importantly, our model can naturally reproduce ring–like and patchy plaque morphologies, as observed experimentally (see Figure 2). Together, these aspects highlight how spatial structure, viral diversity, and innate immunity combine to shape infection outcomes.

**Figure 1:**
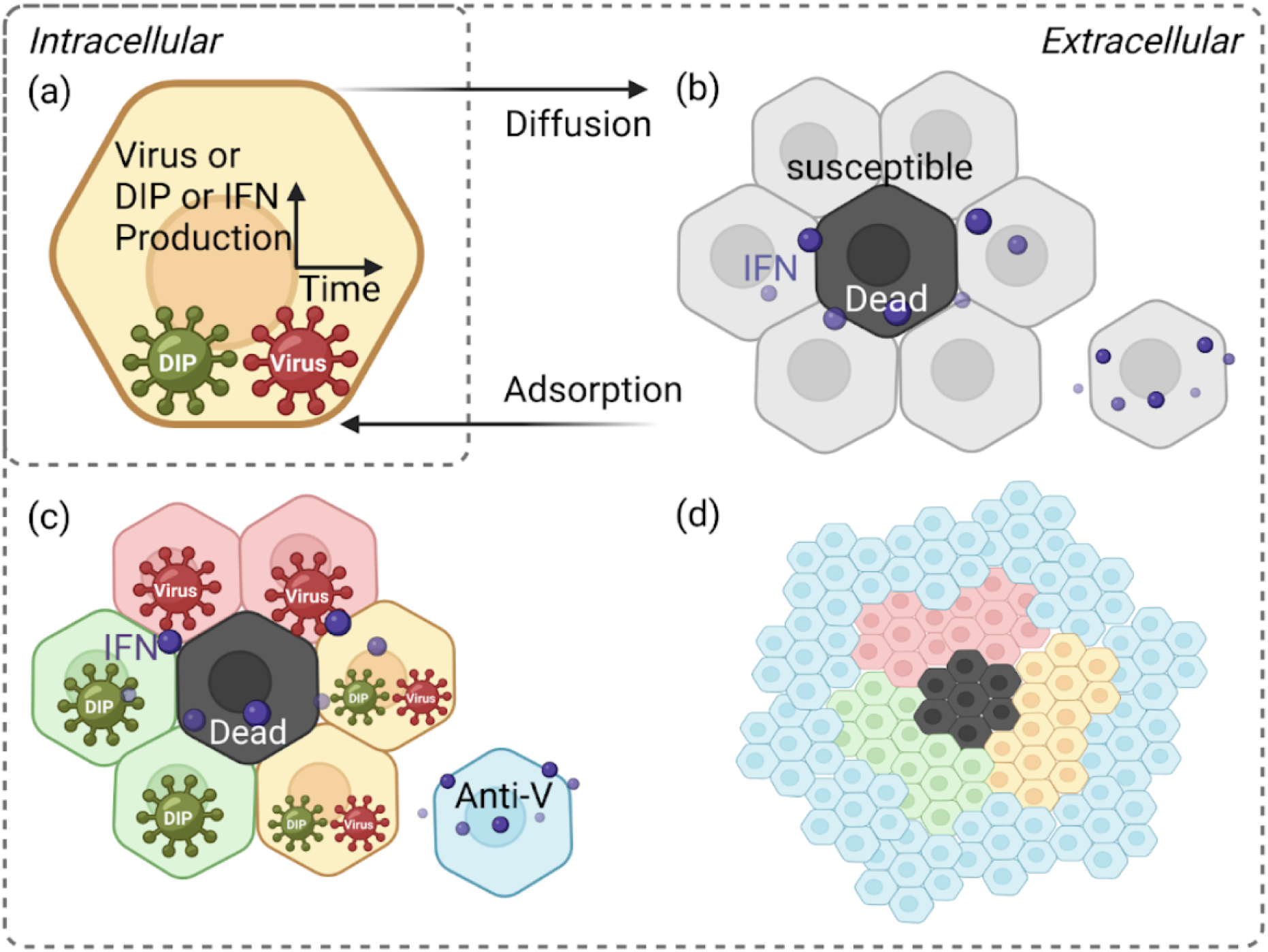
Schematic of the spatially explicit cell–based model. The model represents cell interactions and infection dynamics on a hexagonal lattice, capturing spatial organisation and simulating realistic cell–to–cell spread together with IFN effects. Particles are shown as dark green for DIPs and dark red for viruses. Cell states are colour–coded: light red for virus–infected cells, light green for DIP–infected cells, yellow for cells co–infected by both viruses and DIPs, black for dead cells, blue for antiviral cells, and grey for susceptible cells. (a) Intracellular stage: each infected cell produces viruses, DIPs, and IFN over time. (b) Initial infection: infected cells release viruses and DIPs upon cytolysis, which disperse to neighbouring cells. IFN reduces infection probability and induces antiviral states in uninfected cells. (c) Local spread: viruses infect neighbouring susceptible cells, leading to virus–only, DIP–only, or co–infected states. IFN continues to propagate antiviral effects, preventing further infection of antiviral cells. (d) Antiviral state and recovery: over time, cells switch to an antiviral state, halting spread. Dead cells are replaced through regrowth, restoring the population. Figure adapted from Howat et al. [26].

**Figure 2:**
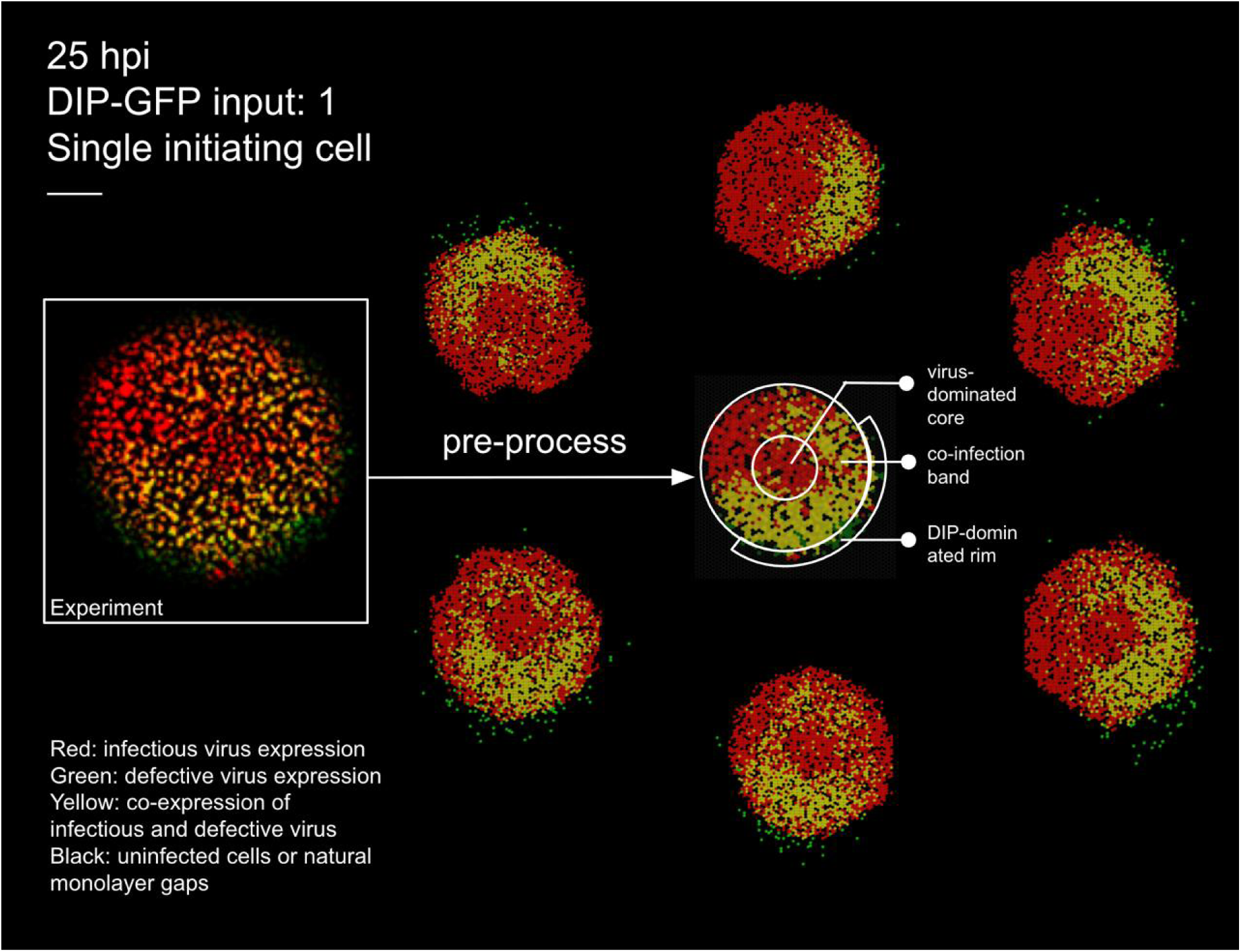
Experimental plaque and matched stochastic simulations at 25 hpi. Left: raw experimental image from [32] with DIP–GFP input of 1 (single initiating cell). Centre: pre–processed experimental image obtained by tessellating into hexagonal cells and applying automated colour classification, which resolves cell states even when colours are too faint to be distinguished by eye. This processed representation is used to count the number of cells in each state for quantitative comparison with simulations. Surrounding six panels: stochastic realisations under matched inoculation and field size, illustrating variability across runs. Colours: black, susceptible; red, virus–infected; green, DIP–infected; yellow, co–infected. Scale bar: 0.5 mm.

These multiple components are important because immune responses are inherently heterogeneous due to the structural complexity, diverse cell populations, and spatially varying microenvironments of human organs. For instance, in the lung, alveolar and bronchial regions exhibit distinct immune activities, with alveoli specialised for gas exchange and bronchi acting as physical barriers to infection [34, 35, 36]. Similarly, in the liver, during hepatitis B infection, the virus may exploit regions of low immune activity near central veins as “safe zones”, evading immune clearance and underscoring the need to include spatial and immune heterogeneity in infection models [37]. While *in vivo* complexity motivates the need for a more biologically refined and spatially granular infection model, *in vitro* systems provide a controlled setting to isolate and model key spatial–immune interactions, helping bridge the gap and guide future experiments.

### Application to IAV

IAV is a highly contagious respiratory pathogen that causes seasonal epidemics and occasionally severe pandemics, providing a well–established system in which to study DIP–mediated modulation of infection [38, 16]. We parameterised key infection and immune timing parameters—such as virus and DIP burst sizes, IFN delay, and IFN decay—using values derived from published IAV infection experiments [39, 40] (these parameter choices are described in detail in the Methods section). With these values, our model captures the interplay between viral diversity and IFN under biologically grounded conditions. During replication, IAV DIPs arise from internal deletions in one of the virus’s eight RNA genome segments due to polymerase template switching. These DelVGs are typically 300–500 nt long, retaining 200 nt from each end of the full–length segment, which ranges from 890 nt (NS) to 2,341 nt (PB1/PB2) in length [41]. They serve as replication templates and are packaged into viruses, with most originating from polymerase–encoding segments [42, 43].

### Motivating questions

How does IFN spatial behaviour change under different diffusion assumptions—instantaneous versus finite–rate—and how does this influence immune protection? What happens if viruses, including DIPs, are not well–mixed, with some jumping across longer distances and generating infection patterns and innate immune responses akin to mucus–mediated dispersal in the respiratory tract? Is a larger DIP output always associated with a stronger IFN response? How do plaque dynamics—such as timing, peak virulence, and infected area—change with or without DIPs or IFN? How can we explain diverse plaque morphologies observed experimentally, including patchy structures, within a unified spatial–immune framework? These questions can be explored using our flexible online RShiny app, which allows users to visualise viral spread, IFN diffusion, and particle movement through dynamic videos; generate time series plots of cell states, virus counts, and IFN levels; and analyse how outcomes vary across different parameter settings (https://shiny-spatial-infection-app-production.up.railway.app/).

## 2 Materials and Methods

Because spatial infection data are crucial for constraining our framework, we first calibrated the model against available experimental datasets. Full datasets that jointly track IFN and DIP dynamics are not yet available, but we can benchmark against published IFN-only and DIP-only experiments to capture complementary aspects of the system and ensure the model reproduces key qualitative features.

### 2.1 Overall model structure

We start with a hexagonal lattice structure to represent the confluent epithelial monolayer, because epithelial cells adhere tightly during *in vitro* growth (Figure 1; [26, 44]). This lattice–based, spatially explicit framework allows us to model individual cell interactions and capture the localised nature of infection, which cannot be approximated by homogeneous mixing or continuum diffusion models [45, 46, 47, 48]. Cell connections are weighted by distance on the lattice, while the infection probabilities associated with these connections are updated hourly to capture the changing dynamics. The model defines seven cell states: susceptible, infected by both DIPs and viruses, infected by DIPs only, infected by viruses only, antiviral, dead, and regrowth (see Figure S1 for the schematic of model execution and explicit model transition rules).

### 2.2 Initial conditions

At the start of a simulation, all cells are susceptible. The standard grid size is 360 × 360 cells, matching the experimental field size in Howat et al. [26], while reduced 50 × 50 grids are used for small–scale tests. Viruses and DIPs are seeded by assigning initial particle counts to randomly selected cells, with *V*_PFU_INITIAL_ = 1 and *D*_PFU_INITIAL_ varying by experimental condition.

### 2.3 Infection probability and IFN response

For A549 cells (which mount a robust IFN response to infection), the probability of infection for both viruses and DIPs is given by

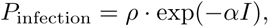

where *ρ* = 0.026 as estimated for influenza A virus *in vitro* plaque assays [26], *I* is the IFN concentration, and *α* = 1.5 is the suppression factor (estimated from the same system). The IFN concentration is updated hourly under either local or global signalling. Guided by experimental observations, we initialised simulations so that DIP-only infected cells secrete more IFN (≈5-fold relative to virus-only cells) and co-infected cells even more (≈10-fold), with these fold changes serving as adjustable starting values [31, 3]. IFN induces an antiviral state in neighbouring cells after a mean delay of *τ* = 12 h (standard deviation 3 h), with a half–life of 3 h [39, 40]. Antiviral cells cannot be infected, creating local barriers to spread. In this setting, the virus is sensitive to the IFN response it provokes.

We implemented two distinct IFN signalling modes: (i) **Finite–range/local signalling**, where IFN spreads diffusively over a limited radius, creating spatial gradients of protection; and (ii) **Global/instantaneous signalling**, where all cells share a uniform IFN concentration at each time step, corresponding to rapid homogenisation in well–mixed environments.

For Vero cells (which lack an IFN response), *P*_infection_ = *ρ* with all IFN–related parameters set to zero. Vero cells therefore remain susceptible throughout infection and never enter the antiviral state [26].

### 2.4 Infection cycle and particle release

Cells infected by viruses or co–infected progress to lysis after an average of 12 h (standard deviation 3 h), releasing progeny. Virus burst size follows a normal distribution with mean 50 PFU per cell, while DIP burst size averages 100 PFU per cell. DIP–only infected cells do not lyse but may convert to the antiviral state. Released particles disperse to neighbouring sites, with local particle density modulating the chance of successful entry.

### 2.5 Cell lysis and regrowth

Lysed cells regenerate when in contact with a neighbouring healthy cell. Regrowth time follows a normal distribution with mean 24 h (standard deviation 6 h). Regrown cells revert to a susceptible state, allowing repeated infection.

In our framework, progeny particles released at lysis can spread in three distinct ways that mirror *in vitro* culture conditions: (i) **Cell–to–cell spread**, which approximates experiments where an agarose or methylcellulose overlay is applied to restrict particles to direct local contacts; (ii) **Finite–range dispersal**, where particles move over several cell diameters, modelling partial diffusion in semi–restricted environments; and (iii) **Unrestricted jumps**, where particles disperse freely across the culture dish without an overlay, as in standard liquid medium, allowing stochastic long–range seeding of secondary plaques.

### 2.6 Feedback dynamics

The cycle of infection, lysis, IFN release, antiviral conversion, cell death, and regrowth generates a feedback loop that drives complex spatial and temporal infection dynamics. This structure provides the foundation for model calibration and experimental comparison described below.

## 3 Results

### 3.1 Model calibration

#### 3.1.1 Calibration with DIP data in the absence of IFN

We first included DIPs and disabled IFN signalling to calibrate our model, following the experimental conditions of Baltes *et al.* [32], who engineered fluorescently labelled DIPs of vesicular stomatitis virus (VSV). Figure 2 compares experimental data with simulation outputs under matched inoculation (DIP–GFP input of 1) and field size.

To enable cell–by–cell quantification, the experimental image at 25 hpi (left) was pre–processed by tessellating the field into hexagonal cells and applying automated colour classification. This procedure identifies infection states (virus only, DIP only, or co–infected) even when colours are too faint to be distinguished by eye, yielding a representation from which the number of cells in each category can be counted. The resulting processed image (centre) forms the basis for quantitative comparison with the surrounding six stochastic replicates generated under a fixed parameter set (see Figure 3 for temporal validation and Methods for details of parameter calibration).

**Figure 3:**
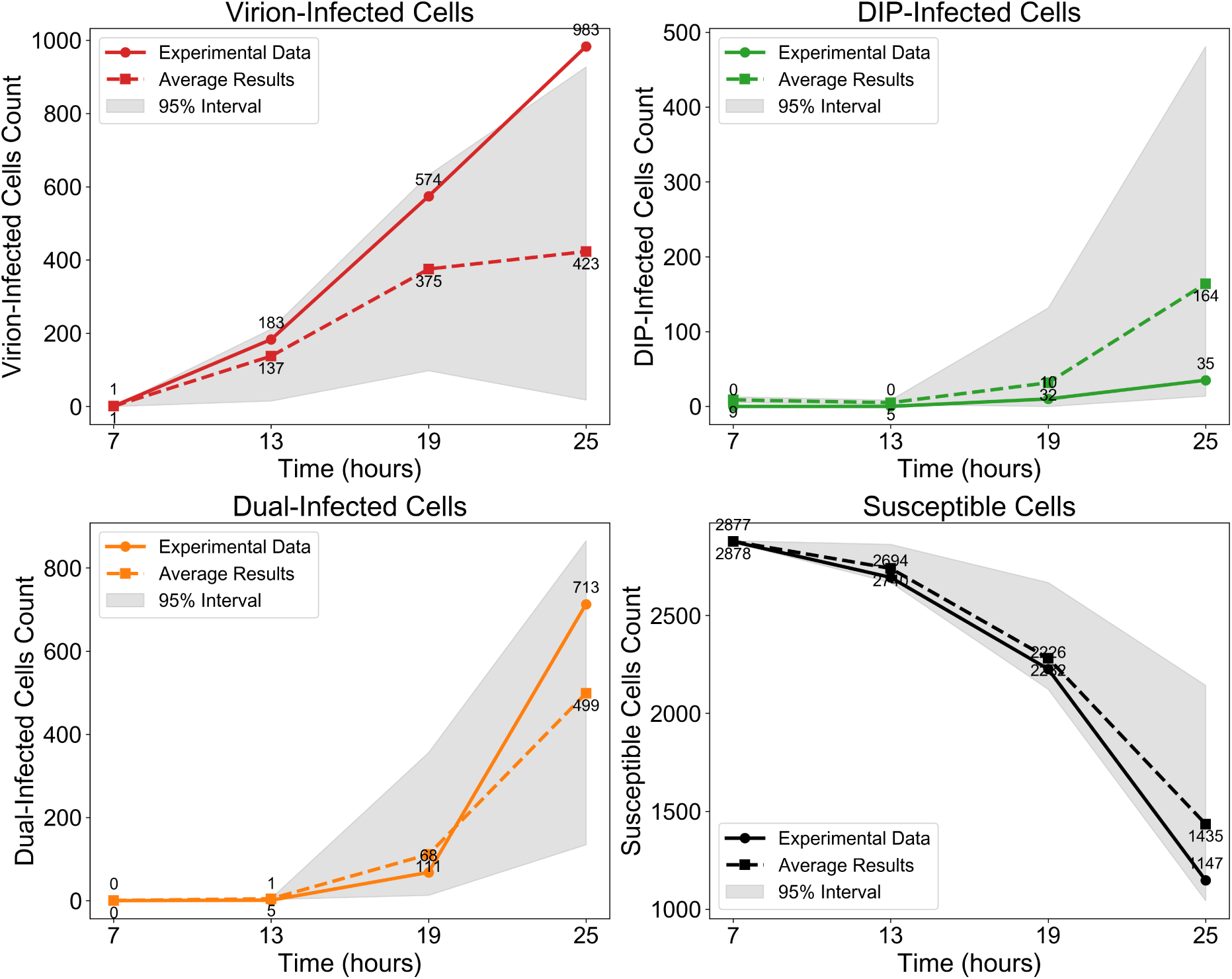
Model validation against experimental data from [32] in the absence of IFN. Cell counts are shown on a logarithmic scale for (top left) virion–infected cells, (top right) DIP–infected cells, (bottom left) co–infected cells, and (bottom right) susceptible cells at four time points (7, 13, 19, and 25 hpi). Solid lines represent experimental data, dashed lines show the mean of stochastic simulations with identical parameters, and grey shading indicates the 95% interval across these simulations.

Parameter values were estimated by scanning plausible ranges for virus and DIP spread (7–35 cells) and burst sizes (50–300 particles per lytic event). Each parameter combination was simulated 30 times, and averages were compared to experimental counts of virus–only, DIP–only, and co–infected cells. The best-fitting set was selected for subsequent analyses.

##### Ring–like and patchy structures emerge in the absence of IFN

Our model reproduces the ring–like and patchy plaque morphologies observed experimentally. These features arise naturally from particle–level stochastic dynamics, without requiring image binarisation or geometric segmentation as in other approaches [33]. Figure 2 illustrates this comparison: the left panel shows the experimental image, the centre panel the pre–processed tessellated representation, and the surrounding six panels representative stochastic simulations.

The central circular region is initially dominated by virus–infected cells (red). Due to the use of maximum–intensity projection, as in the experiment [32], the red signal persists for an extended period, reflecting the long half–life of RFP/GFP even after lysis and the presence of dead cells in the centre. In one sector of the advancing front, DIPs randomly seed a neighbouring region (green). Where these fronts overlap, a circumferential co–infected band appears (yellow).

The band is annular rather than a local patch, because the central red signal persists under maximum–intensity projection even after lysis; the overlap therefore wraps around the circular core. An outer green rim forms because DIPs disperse farther than viruses. This mechanism explains the rings in Figure 2 and is consistent with the concentric–ring analysis and localised enrichment described in the experiment [32].

Beyond morphology, the model also captures the temporal dynamics of infection classes (Figure 3). Although variability across stochastic replicates is wide (grey 95% confidence intervals), the mean trajectories closely follow experimental trends, supporting the model’s ability to reproduce both spatial and temporal aspects of DIP–virus interactions. The main discrepancies arise from stochasticity in whether initial DIP foci encounter virus and establish coinfection. When coinfection occurs early, both viruses and DIPs generate large bursts of progeny and sustain spread; when DIPs fail to meet virus, they quickly fade because they cannot replicate on their own.

#### 3.1.2 Calibration with IFN responses in the absence of DIPs

After calibrating spatial heterogeneity using the Baltes dataset, we next validated IFN–mediated responses using the experiments of Howat *et al.* [26], who analysed Herpes simplex virus 1 (HSV–1) infections in MDBK (IFN–competent) and Vero (IFN–deficient) cells [26, 49, 50, 51, 52]. We set the antiviral delay time to 95 h, consistent with Howat et al. [26], which ensured that our framework captured IFN activation kinetics (Figure 4; see Methods for details of parameter calibration), giving a good agreement with the experimental data.

**Figure 4:**
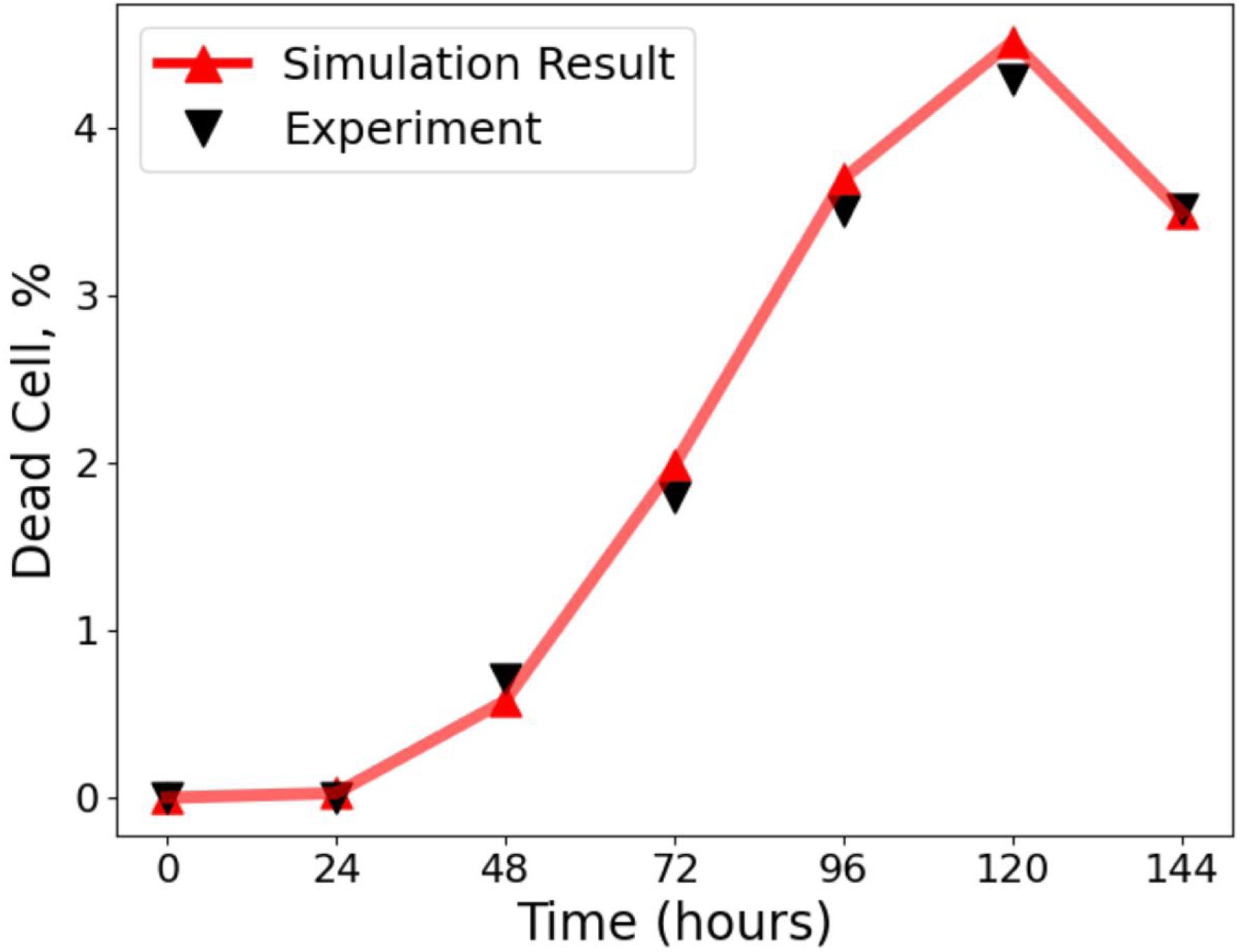
Comparison of model simulations with experimental data from Howat et al. [26]. The percentage of dead cells over time is shown, with simulation outputs (red line, triangles) closely matching experimental measurements (black triangles). Simulations used a time step of 1 h, and the virion infection probability was re–fitted to account for the susceptibility of MDBK cells to HSV infection, giving parameter estimates *ρ* = 0.09 [0.07, 0.11], *τ* = 95 [88, 101] h, and *α* = 1.5 [1.2, 1.8]. Experiments were initiated with an inoculum of 50 pfu, where the pfu represents a statistical average of plaques expected across repeated assays. Each simulation curve represents the mean of 30 independent realisations. This validates the parameterisation of our framework against the dataset of Howat et al. [26]

##### From calibration to IAV simulations

Together, these complementary calibration datasets—VSV with DIPs in the absence of IFN (Baltes *et al.* [32]), and HSV with IFN in the absence of DIPs (Howat *et al.* [26])—provide independent validation of our framework. Although obtained in different viral systems, they establish spatial and immune benchmarks that we used to parameterise subsequent simulations of IAV, the focus of this study.

#### 3.1.3 Dynamics of the full DIP–IFN model

To examine how IFN dynamics shape infection outcomes, we varied three factors in all combinations: IFN response mode, particle movement, and composition. In Figure S2, rows correspond to IFN mode (1 and 4: no IFN; 2 and 5: finite–range; 3 and 6: global/instantaneous), while columns correspond to particle movement (1: cell–to–cell; 2: finite–radius jumps; 3: random long–range jumps). The figure is also divided by composition: the top three rows show viruses with DIPs, whereas the bottom three rows show viruses only. The left panels display temporal changes in cell states and IFN levels, and the right panels show final plaque morphology. These simulations provide a systematic overview of model behaviour and establish a framework for later comparisons (see Figure S2).

### 3.2 Emergent properties of the full model

#### 3.2.1 Intermediate relative DIP yield maximises peak IFN

To investigate how variation in DIP production affects the spatial dynamics of IFN activation, we varied the *relative DIP yield* 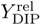, defined as the DIP:virus output ratio upon lysis of a virus–infected cell. In our model, 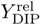 ranged from 2 to 32, corresponding to DIP burst sizes of 100 to 1600 PFU/cell, assuming a fixed virus burst size of 50 PFU/cell based on experimental data and the baseline model [22]. We implemented global IFN signalling and restricted virus dispersal to cell–to–cell spread, using parameter values consistent with previous studies [53, 54, 49]. Simulations were run for 500 hours, with hourly measurements of plaque size, IFN concentration, and infection composition.

We observed a nonmonotonic relationship between 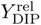 and IFN response, with a clear peak at an intermediate value of 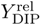, where the IFN concentration reached its maximum (Figure 5, left panel). Beyond this point, further increases in DIP output led to diminished IFN activation. This critical value coincided with the highest number of cells co–infected by both viruses and DIPs, which in our model are the strongest IFN producers—releasing ten times more IFN than virus–only infected cells and twice as much as DIP–only infected cells. At this yield, the virus–only infected population was minimal, while DIP–only infections were widespread. This configuration promoted efficient DIP dissemination, followed by sufficient virus co–infection to generate a large number of double–infected cells.

**Figure 5:**
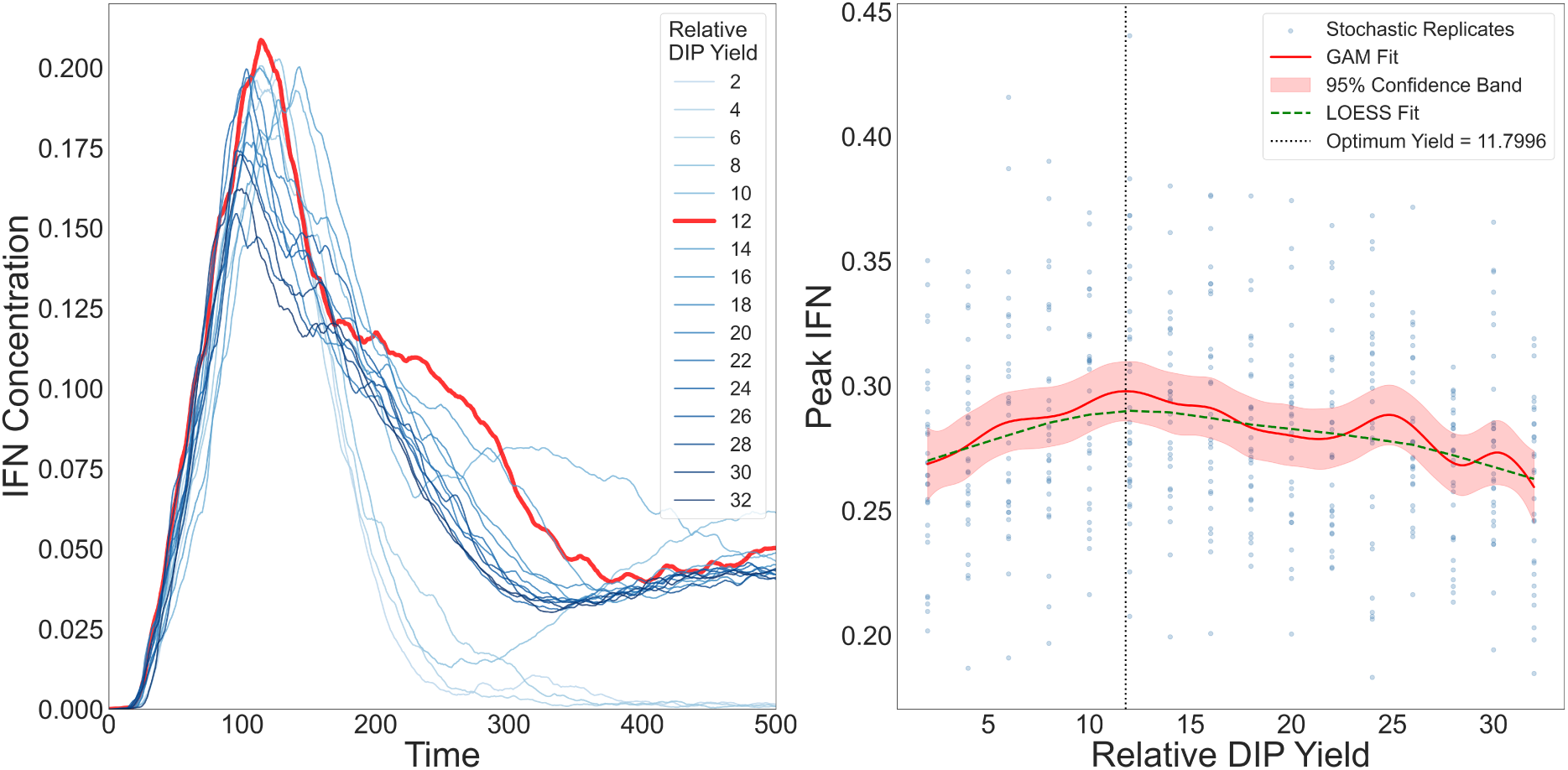
IFN dynamics across values of the *relative DIP yield* 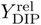. **Left:** IFN time series for increasing 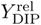 the response rises and then declines, with a maximum at approximately 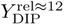 (thick red trace). **Right:** Peak IFN versus 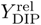 across stochastic replicates. Blue points show replicates; the GAM fit (red, with red 95% confidence band) and the LOESS fit (green, dashed) agree on a non–monotone pattern with an intermediate optimum (vertical dotted line; its exact location shifts with smoothing parameters but always lies at intermediate 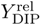).

To make sure that the observed nonmonotone optimum is not a random outcome of stochastic variability, we further analysed replicate simulations using generalized additive models (GAMs) with 95% confidence interval and LOESS smoothing (Figure 5, right panel). Both methods consistently recovered an interior maximum in peak IFN across the explored values of the 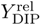. The exact location of this maximum may shift with the spline dimension and smoothness penalty, but the qualitative pattern is robust: peak IFN is highest at intermediate 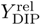, rather than increasing monotonically with 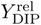.

The observed peak arises from the balance between two opposing regimes (Figure 5):

1. At low 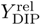, DIP production was insufficient to support widespread DIP–only infections or effective co–infection. Consequently, the number of double–infected cells remained low, and total IFN output stayed limited despite virus spread.
2. At high 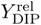, DIPs spread rapidly, initiating early co–infection and triggering premature IFN release. Due to the exponential sensitivity of infection probability *P*_infection_ to IFN levels, both DIPs and viruses quickly lost the ability to infect new cells, curtailing further spread and reducing cumulative IFN production.

Thus, an intermediate peak value of 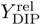 marks the most effective balance between particle production, co–infection timing, and IFN–mediated feedback. Values above or below this point disrupt the coordination between DIP seeding and virus co–infection, ultimately leading to reduced IFN activation.

#### 3.2.2 Minimal fractions of free–jump particles strongly enhance spread

To assess sensitivity to long–range transmission (e.g., mucus–mediated dispersal), we simulated infections with free–jump fractions from 1% to 90% in the absence of IFN; the remaining transmission occurred by cell–to–cell spread. As shown in Figure S3 (top left), even 1% free jumps accelerated plaque expansion and altered morphology, with satellite plaques visible by *t* = 150 h.

Under global IFN signalling, free jumps had little effect because uniform antiviral activation removed any advantage of escaping local immune zones (Figure S3). In contrast, under local IFN signalling, even 1% free jumps had a strong impact.

Figure 6 shows these outcomes in detail. Despite an earlier and higher antiviral fraction with 1% free jumps (blue curves) than with no jumps (cyan), the dead–cell fraction (red curves) spikes much earlier and reaches peaks comparable to the no–jump IFN10 condition (orange). The reason is the response delay *τ*: free jumps establish foci beyond the contemporaneous IFN front, allowing damage to accumulate before local antiviral containment arrives.

**Figure 6:**
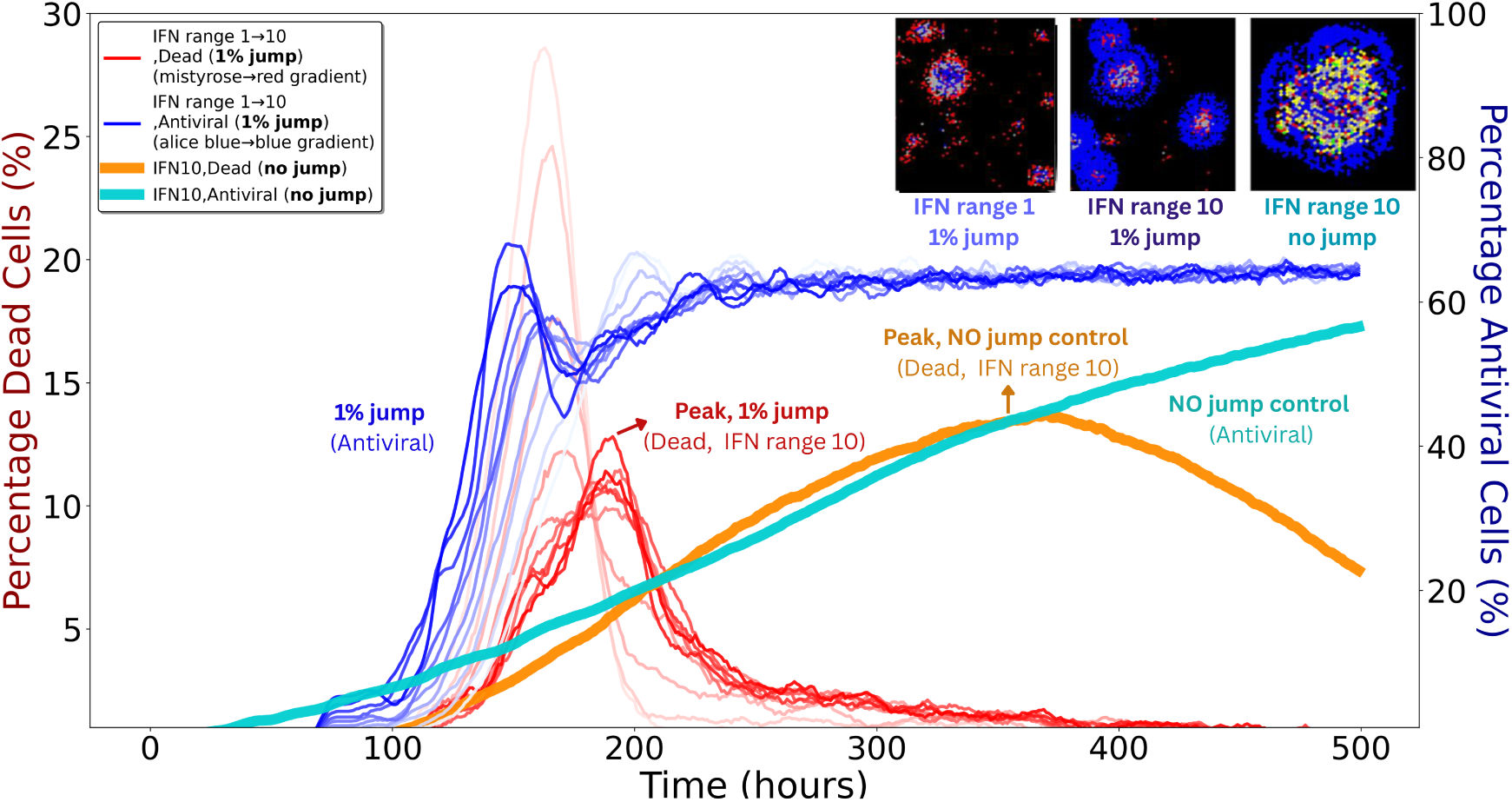
IFN range and spreading mode jointly shape plaque dynamics. Thin red and blue curves denote simulations with 1% free–jump transmission for both viruses and DIPs (remaining 99% cell–to–cell). Red gradients (left *y*–axis) show dead–cell fractions for IFN ranges 1–10, where free jumps generate earlier and higher peaks (arrow, “Peak, 1% jump”). Blue gradients (right *y*–axis) show antiviral–cell fractions for the same simulations, rising earlier and to higher levels than the no–jump control (labelled “1% jump, Antiviral”). Thick orange (dead) and thick cyan (antiviral) curves indicate IFN10 controls without free jumps; the orange curve peaks later (arrow, “Peak, no jump control”) and at similar magnitude to the red curves, while the cyan curve shows a slower antiviral rise (labelled “No jump control”). The three insets (top right) show spatial patterns at *t* = 120 h: left, IFN1 with 1% free jumps yields scattered secondary plaques; middle, IFN10 with 1% free jumps forms larger but more contained plaques; right, IFN10 with no free jumps produces a single compact plaque. Inset colours: black, susceptible/uninfected; grey, dead; red, virus–infected; green, DIP–infected; yellow, co–infected; blue, antiviral.

Increasing the IFN range provides stronger control: as the range grows, dead–cell peaks (red, orange) are reduced and delayed. The antiviral trajectories (blue, cyan), however, remain similar across ranges because their timing is largely set by *τ*. Thus, outcomes depend mainly on how many new foci are seeded outside the current IFN domain rather than on modest differences in IFN radius. Jumps that land beyond the front are particularly consequential, and DIPs tend to jump farther than viruses, increasing the likelihood of such events. Each new focus is rapidly encircled by antiviral cells and then declines, but repeated seeding during the delay accumulates damage.

## 4 Discussion

To our knowledge, there is no spatially explicit *in vitro* modelling framework that integrates DIPs and IFN responses. This work complements earlier lines of work: cellular automaton studies examined coinfection and plaque morphology in DIP–rich settings [9, 27, 29] but did not include IFN signalling, while ODE–based IFN models [25, 22] captured timing and magnitude of responses but abstracted away spatial structure. We use this agent–based approach to bring these perspectives together by embedding DIP–IFN interactions in a spatial, stochastic framework. Our simulations are not intended to deliver precise predictions, but rather to generate mechanistic insights that can guide experimental design and future model refinement.

Building on this framework, we examined how spatial spread depends on IFN signalling and particle movement. We considered combinations of IFN spread (none, finite–range/local, global/instantaneous) and dispersal modes (cell–to–cell, finite–radius jumps, rare long–range jumps), under conditions with and without DIPs (Figure S2).

As an initial validation, we calibrated the model to the conditions of *Baltes et al.* [32] to test qualitative consistency with the DIP–only dataset lacking IFN and to establish stochastic variability as a baseline for subsequent analyses. We also confirmed the IFN response dynamics by calibrating the model to the *Howat et al.* [26] dataset, which captures antiviral activation in MDBK and Vero cells under defined infection conditions. Together, these complementary datasets ensure that both spatial and immune aspects of the model are quantitatively grounded.

This experiment provides valuable insight into spatial spread in the presence of DIPs, but several key quantitative features are not available, especially data on the spatial extent of dead cells. Despite these constraints, our model reproduces the qualitative patterns: a virus–positive centre with ring–wise shifts in reporter dominance, patchy gene expression, and localised regions of DIP or coinfection enrichment, consistent with the concentric–ring analysis (Figure 2). Unlike *Liang et al.* [33], who analysed the same dataset with a PDE framework by dividing plaques into high– and low–density sectors, our agent–based model generates ring–like structures without imposed segmentation (Figure 2). This naturally patchy outcome arises because (i) we apply maximum–intensity projection, as in the experiment, so the initial red signal from lysed cells persists and maintains a virus–positive centre even after clearance, and (ii) in these experiments, DIPs disperse more broadly and stochastically in direction, leading virus–infected regions to remain red while later DIP infection produces more distant and patchy patterns. To help explore and interpret additional experimental phenomena, we provide a flexible RShiny app as an online platform designed to support such investigations.

We next analysed how IFN activation is regulated by relative DIP yield. In our implementation, coinfected cells were the strongest IFN producers (ten–fold above virus–only and roughly two–fold above DIP–only), and maximal IFN arose at an intermediate DIP yield that favoured broad DIP seeding followed by timely virus coinfection (Figure 5). At low DIP yield, coinfection was scarce and IFN remained modest; at high yield, early IFN curtailed further spread of both DIPs and viruses, limiting the cumulative number of secreting cells. Although our model represents within–cell replication indirectly, this pattern is consistent with reports that early replication kinetics shape the likelihood and timing of IFN induction [55]. We emphasise that yield here is a tissue–level control parameter rather than a direct surrogate for intracellular replication rate.

A notable feature of these analyses is spatial dispersal. Under local IFN signalling, finite–rate IFN was sufficient to halt infection when transmission was strictly local. However, introducing even a small fraction (~ 1%) of free–jump events allowed viruses and DIPs to escape IFN–dense zones and establish secondary plaques by seeding foci beyond the contemporaneous IFN front (Figure 6). These free jumps—potentially reflecting in vivo processes such as mucus–mediated transport or epithelial disruption—led to earlier and higher peaks in dead–cell fractions because IFN induction has a response delay *τ*, during which damage accumulates before local containment becomes effective. Increasing the IFN interaction range improved control by reducing the fraction of space left unprotected at any instant, but the timing of antiviral activation changed little, being largely set by *τ*. By contrast, under global or instantaneous IFN, the advantage of escaping local immune zones was lost, and free jumps had minimal impact. Although the precise dispersal behaviour of respiratory viruses *in vivo* remains uncertain, long–range, spatially discontinuous spread has been documented in murine models, such as inter–lobar transitions following low–dose inoculation [35]. This resemblance echoes the small–world effect, where the addition of just a few long–range connections can shorten path lengths in otherwise locally connected networks [56].

Finally, our results highlight a trade–off in IFN induction by DIPs. Too few DIPs allow wild–type virus to dominate, while too many suppress viral replication before IFN is strongly activated. At intermediate levels, IFN output is maximised, but the trade–off is not straightforward: strong local responses can be bypassed if viruses disperse by rare long–range jumps. Thus, IFN operates effectively only within the spatial domain it reaches, and additional IFN cannot prevent escape once particles move beyond that range. Together, these findings underscore how DIPs enhance protection by balancing containment and escape, suggesting design principles for therapies that aim to exploit or mimic DIP–induced immunity. More broadly, the balance we observe may also shape virus evolution, as the interaction between DIPs, IFN, and dispersal determines which viral strategies can persist.

Notably, beyond IAV, our model is flexible and adaptable to other pathogens, as DIPs are widely observed across viruses such as measles [57], COVID–19 [58], dengue [59], HBV [60], and HIV [61], etc.

### 4.1 Caveats and Model Limitations

The experimental dataset from *Baltes et al.* [32], which includes DIPs but no IFN, provides a useful qualitative benchmark but omits key details such as the spatial extent and timing of cell death, natural monolayer gaps, and particle spreading ranges, all of which affect outcomes but cannot be fitted. Only four time points were imaged, further limiting temporal resolution. In Baltes et al., Fig. 3b, separate red (virus) and green (DIP) channels were shown alongside merged images, but our attempts to digitally recombine the channels did not fully reproduce the published merge, likely due to resolution or processing differences not described in detail. This does not affect qualitative interpretation.

The same dataset was also analysed by *Liang et al.* [33], who noted that reporter intensity is qualitative rather than quantitative. Thus, the images serve as indicators of viral gene expression but do not allow precise parameter calibration. While our simulations reproduce the main experimental features, the absence of dynamic measurements constrains time–resolved predictions. To address this, we also drew on data from *Howat et al.* [26], which include IFN but no DIPs, providing complementary context. As IFN responses can strongly alter spatial spreading dynamics, future studies should consider including clear imaging or other quantitative measurements of IFN diffusion rates and antiviral state transition times in DIP–infected systems, to better resolve DIP–IFN interactions.

A further complication is that IFN production is not a continuous process. Cells do not sustain secretion indefinitely; instead, expression is modulated by transcriptional feedback and other regulatory mechanisms [62]. Our current model does not include such downregulation, which may be important for understanding immune exhaustion and cytokine storm phenomena. In reality, IFN transcription can be switched off through repression, viral antagonism, or intrinsic negative feedback involving SOCS proteins, USP18, and PIAS1, generating temporal heterogeneity even among similarly infected cells. Immune signalling is further shaped by adaptive responses: CD8^+^ T cells and B cells modulate IFN pathways, and their delayed engagement can influence plaque morphology and clearance. These layers of regulation, supported by recent studies [63], suggest that integrating transcriptional and cellular feedback into future models will improve biological realism and link innate with adaptive dynamics.

### 4.2 Future Directions

Future extensions of this model could further clarify how viral and immune processes interact. One improvement would be to introduce stochastic variation in IFN production across cells, capturing the heterogeneity observed in single–cell studies [64]. Another direction is to connect in vivo tissue–level dynamics to population–level spread using multi–scale modelling, which may clarify how DIPs shape transmission and immune control. While our study relies on stochastic simulations, their complexity can obscure mechanistic understanding. Analytical approximations or simplified models may help disentangle the nonlinear feedbacks underlying IFN responses.

From an experimental perspective, future studies should focus on quantifying IFN diffusion and anti-viral state transition kinetics in cell culture systems coinfected with standard virus and DIPs. Although some *in vivo* data on DIP spreading are available [12], they typically provide only a few time points or lack spatial resolution. Time–lapse imaging combined with single–cell RNA sequencing could yield crucial data for refining model parameters, help explain experimental phenomena such as comet–like spread[65], and guide the design of future experiments. In addition, co–culture experiments with DIP–producing and non–DIP–producing viral strains would help validate the predicted effects of DIPs on infection dynamics and IFN responses.

## Declarations

### Authors contributions

Y.L.: conceptualization, methodology, software, validation, formal analysis, writing – original draft, visualization, writing – review and editing; B.T.G.: conceptualization, methodology, supervision, writing – review and editing; A.J.W.t.V.: methodology, software, validation, formal analysis, supervision, writing – review and editing; B.F.N.: validation, software, formal analysis, writing – review and editing; S.A.L.: writing – review and editing.

All authors gave final approval for publication and agreed to be held accountable for the work performed therein.

### Funding

This work was supported by a Princeton Catalysis Initiative award (to B.T.G. and A.J.W.t.V.); Princeton Precision Health (to B.T.G.); the National Institutes of Health (NIH) grant DP2 AI175474 (to A.J.W.t.V.); the Carlsberg Foundation (grants CF23-0173 and CF24-1337 to B.F.N.); the National Cancer Institute, NIH, under Prime Contract No. 75N91019D00024, Task Order No. 75N91023F00016 (the content of this publication does not necessarily reflect the views or policies of the Department of Health and Human Services, nor does mention of trade names, commercial products or organizations imply endorsement by the U.S. Government); and the National Science Foundation (DMS-2327711, Collaborative Research: IHBEM: Data-driven multimodal methods for behavior-based epidemiological modeling).

### Data and code availability

All simulation codes, analysis scripts, and data supporting the results of this study are openly available at: https://github.com/yimei-li/spatial-dynamics.

### Ethics statement

This study did not involve human participants or animals. Ethical approval was not required.

### Competing interests

The authors declare no competing interests.

### Use of Artificial Intelligence (AI)-assisted technologies

AI-assistance was used for grammar correction and completing lines in our code. All AI-produced code information verified by Yimei Li.

## Supporting Information

**Figure S1:**
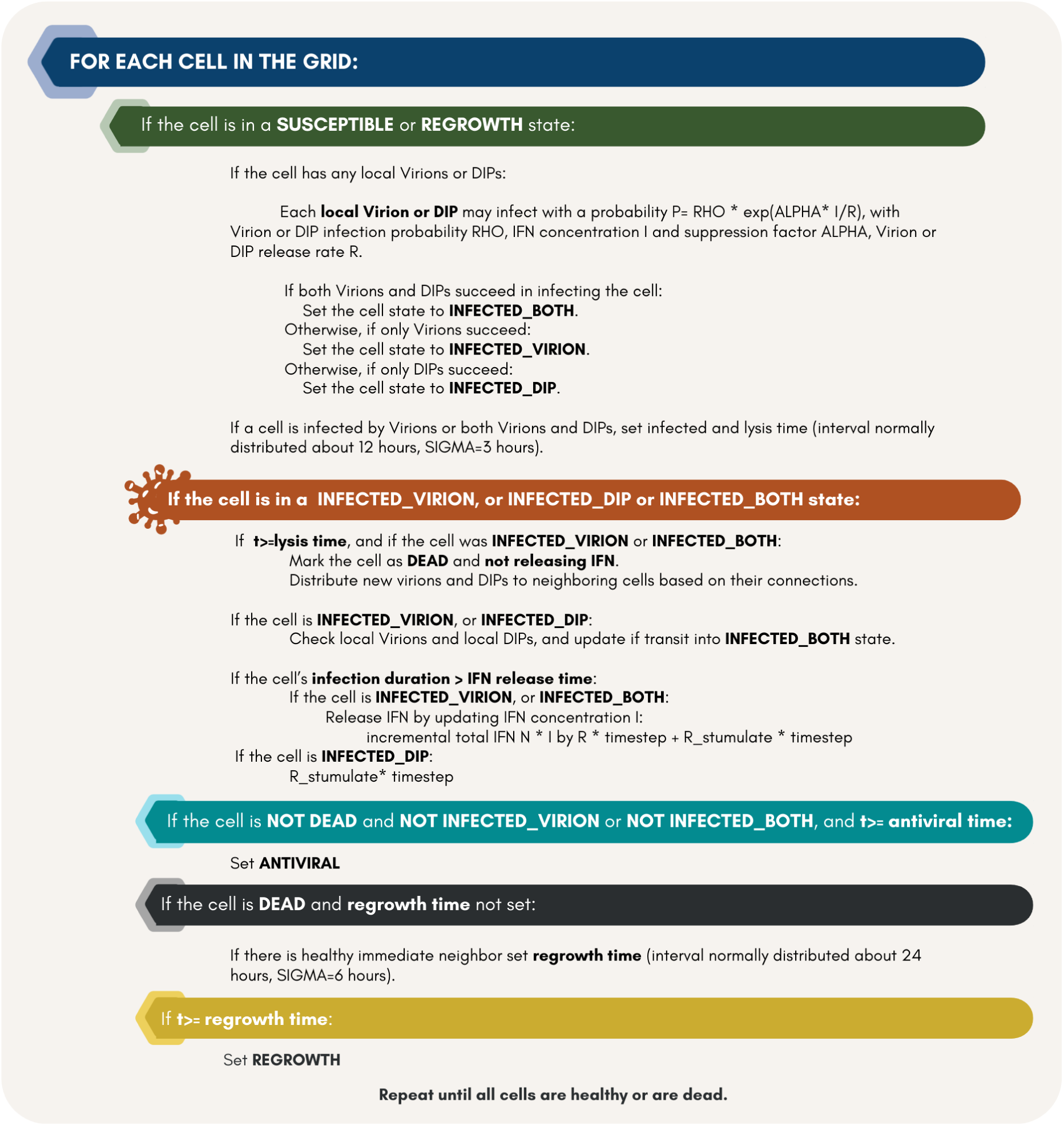
Model execution flowchart and explicit transition rules. The simulation begins by setting up a hexagonal grid where all cells are initially susceptible. Viruses and DIPs are introduced based on the inoculum size. Infection dynamics unfold through stochastic transitions: susceptible and regrowth cells become infected based on local particle and IFN levels. Infected cells undergo lysis, release particles and IFN, and die. Dead cells regrow if healthy neighbours are present. The model continues until termination criteria are met. Output includes time–resolved data and spatial maps. Animations available at: https://shiny-spatial-infection-app-production.up.railway.app/.

**Figure S2:**
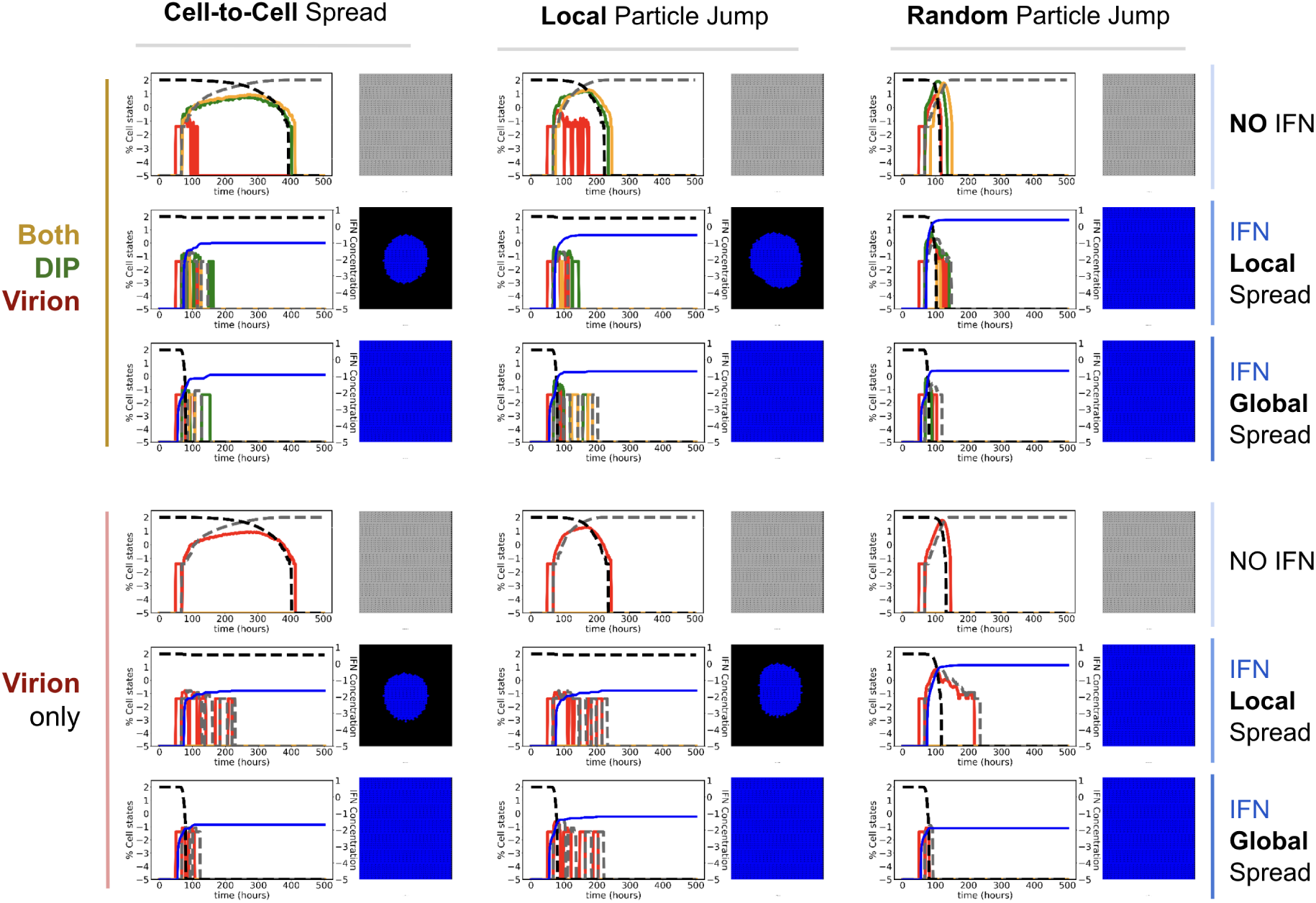
Dynamics of spread under varying particle–jump behaviours and IFN conditions. Outcomes are shown for scenarios with both DIPs and viruses (top three rows) and viruses only (bottom three rows), under three movement modes: cell–to–cell (left column), finite–radius jumps (middle column), and random jumps (right column). Each row corresponds to a different IFN mode: no IFN, finite–range IFN spread, and global/instantaneous IFN spread. **Left plot**: temporal dynamics. Both *y*–axes use a log_10_ scale: the left *y*–axis shows the percentage of cells in each state—**virus–only infected**, **DIP–only infected**, **co–infected**, **regrowth**, and **dead**; the right *y*–axis shows the **global IFN concentration** per cell. **Right plot**: the final spatial distribution of states: **virus–only**, **DIP–only**, **co–infected**, **antiviral**, **susceptible**, and **dead**.

**Figure S3:**
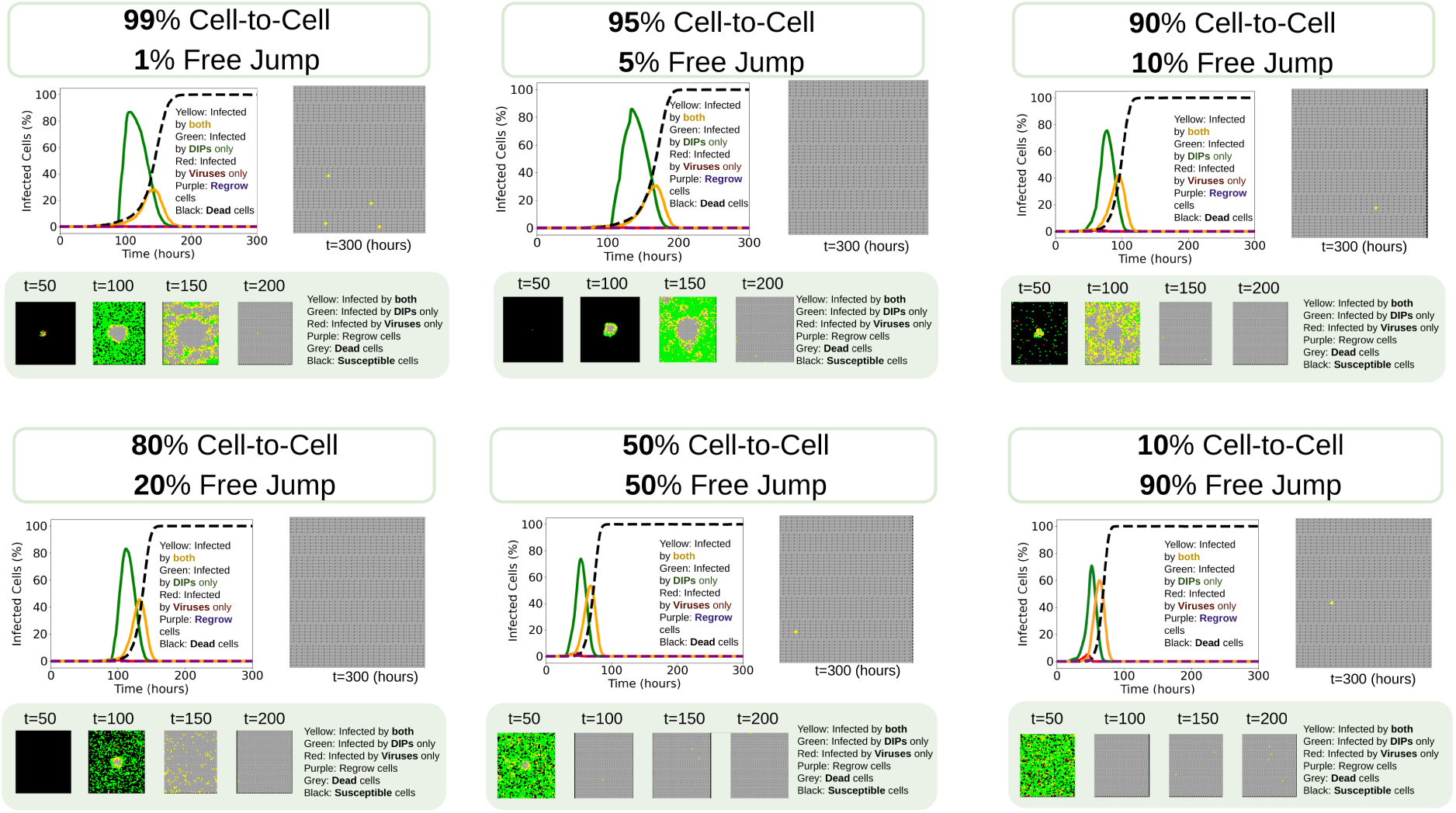
Heterogeneity in infection dynamics and plaque morphology under varying proportions of cell–to–cell and free–jump spread in Vero cells. Left panels: The line plots illustrate the percentage of infected cells over time, including viruses–only infections (red), DIPs–only infections (green), coinfections by both (yellow), regrowth cells (purple), and the dashed line represents dead cells (black). Right panels: The final plaque morphology at the simulation endpoint (t = 300 hours), highlighting the spatial distribution of cell states. At this time point, the plaques (grey) have nearly reached full coverage. Bottom panels: Plaque development over time at t = 50, 100, 150, and 200 hours, showing the progression of infection. Colours in the right and bottom panels represent different cell states: infected by both DIPs and viruses (yellow), infected by DIPs only (green), infected by viruses only (red), antiviral state (blue), dead cells (grey), uninfected/susceptible cells (black), and regrowth cells (purple).

**Figure S4:**
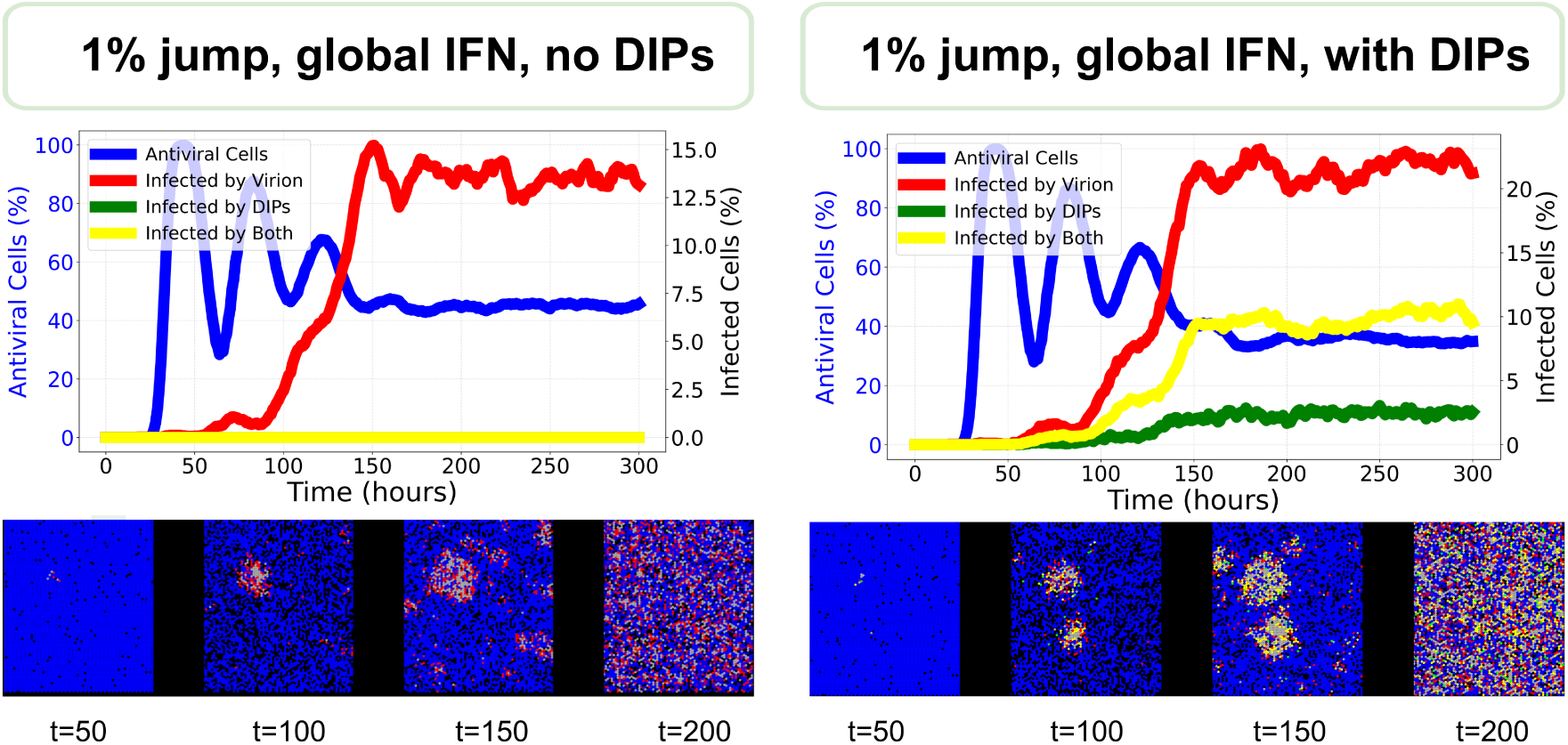
Comparison of plaque dynamics under global IFN with 1% free–jump transmission, in the absence (left) and presence (right) of DIPs. Top panels: time courses showing percentages of antiviral cells (blue, left *y*–axis), virus–infected cells (red), DIP–infected cells (green), and co–infected cells (yellow) (all right *y*–axis). Bottom panels: representative spatial snapshots at *t* = 50, 100, 150, and 200 hours, illustrating how DIPs promote co–infection (yellow) and more diffuse plaque morphology despite similar levels of antiviral activation. Colour code: black, susceptible/uninfected; grey, dead; red, virus–infected; green, DIP–infected; yellow, co–infected; blue, antiviral.

## Notes

https://github.com/yimei-li/spatial-dynamics

https://shiny-spatial-infection-app-production.up.railway.app/

### Competing Interest Statement

The authors have declared no competing interest.

### Summary of Updates

In this revised version, we have corrected incomplete and erroneous references that had inadvertently been added to the original submission.

